# Multiple expression assessments of ACE2 and TMPRSS2 SARS-CoV-2 entry molecules in the urinary tract and their associations with clinical manifestations of COVID-19

**DOI:** 10.1101/2020.05.08.083618

**Authors:** Xiaohan Ren, Xiyi Wei, Guangyao Li, Shancheng Ren, Xinglin Chen, Tongtong Zhang, Xu Zhang, Zhongwen Lu, Zebing You, Shangqian Wang, Chao Qin, Ninghong Song, Zengjun Wang

## Abstract

**Background:** Since December 2019, the novel coronavirus, severe acute respiratory syndrome coronavirus 2 (SARS-CoV-2), first spread quickly in Wuhan, China, then globally. From previously published evidence, ACE2 and TMPRSS2, are both pivotal entry molecules that enable cellular infection by SARS-CoV-2. Meanwhile, increased expression of pro-inflammatory cytokines, or a “cytokine storm,” is associated with multiple organ dysfunction syndrome that is often observed in critically ill patients.

**Methods:** We investigated the expression pattern of ACE2 and TMPRSS2 in major organs in the human body, especially under specific disease conditions. Multiple sequence alignment of ACE2 in different species was used to explain animal susceptibility. Moreover, the cell-specific expression patterns of ACE2 and cytokine receptors in the urinary tract were assessed using single-cell RNA sequencing (scRNA-seq). Additional biological relevance was determined through Gene Set Enrichment Analysis (GSEA) using an ACE2 specific signature.

**Results:** Our results revealed that ACE2 and TMPRSS2 were highly expressed in genitourinary organs. ACE2 was highly and significantly expressed in the kidney among individuals with chronic kidney diseases or diabetic nephropathy. In single cells, ACE2 was primarily enriched in gametocytes in the testis, and renal proximal tubules. The receptors for pro-inflammatory cytokines, especially IL6ST, were remarkably concentrated in endothelial cells, macrophages, and spermatogonial stem cells in the testis, and renal endothelial cells, which suggested the occurrence of alternative damaging mechanisms via autoimmune attacks.

**Conclusions:** This study provided new insights into the pathogenicity mechanisms of SARS-CoV-2 that underlie the clinical manifestations observed in the human testis and kidney. These observations might substantially facilitate the development of effective treatments for this rapidly spreading disease.

## Introduction

With the appearance of SARS-CoV-2 in Wuhan, China, it became a great challenge globally for health authorities to contain the coronavirus disease 2019 (COVID-19) epidemic. On March 11, 2020, with many infected people reported in numerous countries, COVID-19 was declared to be a pandemic by the World Health Organization (WHO)^1^. Two previous separate outbreaks of coronaviruses (severe acute respiratory syndrome, SARS-CoV, and Middle East respiratory syndrome, MERS-CoV) emerged in specific regions of the world before the appearance of SARS-CoV-2^2,3^. SARS-CoV-2 is remarkably similar to SARS-CoV with a 76% amino acid sequence identity and the same receptor, angiotensin-converting enzyme 2 (ACE2). Concomitantly, TMPRSS2, the serine protease for virus Spike (S) protein priming, also was considered indispensable for cell entry by SARS-CoV-2. In other words, ACE2 and TMPRSS2 are proven key molecules for cell entry by SARS-CoV-2^4^

The dominant clinical signs of COVID-19 are fever and cough, which resemble common viral pneumonia. However, in seriously affected individuals, acute respiratory distress syndrome (ARDS) is the most severe pulmonary complication that is triggered by a cytokine storm and results in a high mortality rate^5^. However, in addition to the central symptoms that affect the respiratory system, the possibility of COVID-19 damaging the digestive and urogenital tracts has been reported, possibly a result of high viral loads and the biological functions of ACE2^6^. Moreover, the amplified pro-inflammatory cytokines could spill over into the circulation and induce a “cytokine storm,” leading to multiple organ dysfunction syndrome in critically ill patients^7^. Recent studies have reported damage to the urinary tract in patients infected with COVID-19. A retrospective study by Huang et al. reported the outcomes of 41 COVID-19 patients in Wuhan and stated that three patients developed complicated acute kidney injury (AKI), and all three needed ICU care^8^. Another retrospective, multi-center cohort study demonstrated the epidemiological and clinical characteristics of 191 COVID-19 patients, of whom 27 individuals were reported to have AKI, and ten were treated with continuous renal replacement therapy (CRRT)^9^. A brief review of existing papers that reported urology system impairment associated with COVID-19 infection is shown in ***Table 1***^8-19^.

**Table 1.** The brief review of existing paper with genitourinary system impairment.

| Journal | Author(Year) | Cases |  |  | Event |
| --- | --- | --- | --- | --- | --- |
|  |  | Total | Male | Female |  |
| Lancet | Huang, C., et al (2020)<sup>8</sup> | 41 | 30 | 11 | AKI: 3; CRRT: 3 |
| JAMA | D, W., et al. (2020)<sup>10</sup> | 138 | 75 | 63 | AKI: 3; CRRT: 2 |
| Lancet | F, Z., et al. (2020)<sup>9</sup> | 191 | 119 | 72 | AKI: 27; CRRT:10 |
| Intensive Care Medicine | Ruan, Q., et al. (2020)<sup>11</sup> | 150 | 102 | 48 | AKI:21; CRRT: 5 |
| BMJ | T, C., et al. (2020)<sup>12</sup> | 274 | 171 | 103 | AKI: 28; CRRT: 3 |
| NEJM | WJ, G., et al. (2020)<sup>13</sup> | 1099 | 637 | 459 | AKI: 5; CRRT: 9 |
| Intensive Care Med | WJ, T., et al. (2020)<sup>14</sup> | 174 | 79 | 95 | AKI: 19; CRRT: 7 |
| Intensive Care Med | J, C., et al. (2020)<sup>15</sup> | 102 | 53 | 49 | AKI: 8; CRRT :4 |
| Fertility and Sterility | Pan, F., et al. (2020)<sup>16</sup> | 34 | 34 | 0 | Scrotal discomfort: 19 |
| JAMA | Richardson, S., et al. (2020)<sup>17</sup> | 5700 | 3437 | 2263 | AKI: 523; CRRT: 81 |
| J Am Soc Nephrol | Pei, G., et al. (2020)<sup>18</sup> | 333 | 182 | 151 | AKI: 35 |
| JAMA Netw Open | Li, D., et al (2020)<sup>19</sup> | 38 | 38 | 0 | SARS-CoV-2 positive in semen: 6 |

**Abbreviations:** AKI, acute kidney injury.

Nevertheless, substantial gaps remain in our knowledge of the biology of COVID-19. In this study, we performed an in-depth investigation into the underlying mechanisms of genitourinary system injury caused by COVID-19. We conducted a series of bulk and single-cell bioinformatics analyses based on the evaluation of multi-access and multiple databases. Our study provides a more detailed and comprehensive perspective of the pathogenicity mechanisms of SARS-CoV-2 infection in the human testis and kidney.

## Methods

### Clinical and epidemiological information

The clinical and epidemiological data for COVID-19 were downloaded from an open-access and continuously updated resource (https://tinyurl.com/s6gsq5y) on April 22, 2020^20^. The visualization map of known COVID-19 cases, worldwide, was available on the website https://coronavirus.jhu.edu/map.html (John Hopkins University).

### Tissue resources

The expression values of ACE2 and TMPRSS2 mRNA and protein in normal tissues were obtained from the following databases, Human Protein Atlas (HPA), The Functional Annotation of The Mammalian Genome (FANTOM5), and The Genotype-Tissue Expression (GTEx). Briefly, the HPA database supplied a map of the transcriptome and proteome profiles across 32 types of tissues and approximately 40 different cell types in the human body^21^. The FANTOM5 project, an international collaborative effort initiated by RIKEN, collected transcriptome data based on more than three thousand human and mouse samples, which involved all major organs in the human body and over 200 cancer cell lines^22^. GTEx provided mRNA expression data from 54 separate normal tissue sites across nearly 1,000 individuals^23^.

The mRNA expression data of ACE2 and TMPRSS2 from 33 cancer types and normal counterparts also was obtained in TCGA using the author’s R codes. Through the Gene Expression Omnibus (GEO) databases, including major human organs, GSE3526^24^, and GSE8124^25^ were chosen to detect ACE2 and TMPRSS2 mRNA expression values. Simultaneously, to explore the different ACE2 and TMPRSS2 expression levels in different people, and using sample counts that were greater than 20, the following genes were selected, GSE113439 (lung tissue; pulmonary arterial hypertension vs. control), GSE66494 (kidney tissue; chronic kidney disease vs. control)^26^, GSE96804 (kidney tissue; diabetic nephropathy vs. control)^27^, and GSE10006 (airway epithelium; smoker vs. non-smoker)^28^.

### Multiple sequence alignment - ACE2

The reported ACE2 gene sequences from different species were downloaded from the National Center for Biotechnology Information (NCBI, https://www.ncbi.nlm.nih.gov/), which included humans (Homo sapiens), domestic chickens (Gallus gallus), waterfowl (Tadorna cana), cats (Felis catus), domestic ferrets (Mustela Putorius furo), wild boars (Sus scrofa), dogs (Canis lupus familiar), raccoons (Procyon lotor), masked palm civets (Paguma larvata), tigers (Panthera tigris altaica), domestic cattle (Bos taurus taurus), and raccoon dogs (Nyctereutes procyonoides). Using MEGA X software, we analyzed differences in gene structure and identified molecular markers via sequence alignment. MEGA X, with the kimura-2 parameter (K2P) model, was used to calculate the genetic distance within species and build a Neighbor-Joining (NJ) tree^29^.

### Single-cell RNA sequencing (RNA-seq) analysis

We used Kidney Interactive Transcriptomics (http://humphreyslab.com/SingleCell/) to visualize the distribution of ACE2 in each cell cluster of kidney epithelia and entire kidneys^30,31^. Two adult testis tissues (Adult 1, unfractionated and Adult 2, unfractionated) of GSE124263 were used to visualize the ACE2 expression pattern in testis cell subgroups^32^. Human Cell Landscape (http://bis.zju.edu.cn/HCL/index.html) was used to visualize the expression patterns for ACE2 and multiple cytokine receptors (CSF1R, CSF2RA, CSF2RB, FGFR2, IFNGR1, IFNGR2, IL6R, IL6ST, IL7R, IL10RA, IL10RB, IL17RA, and NGFR) in kidney and testis single cells^33^.

### Gene-set enrichment analysis for the ACE2-specific signature

GSEA (version 4.0.3) was performed in specific cells to explore further the biological processes associated with ACE2-positive cells. The gene set was C5.BP.v7.1.

## Results

### Clinical and epidemiological features of COVID-19 at diagnosis

***Table S1*** shows the clinical features of 1,911 patients infected with SARS-CoV-2. At the onset of infection, the most common symptom was fever (1,002 of the 1,911 patients, 52.4%), followed by cough (566 of the 1,911 patients, 29.6%). The visualization of COVID-19 epidemiological information is shown in ***Figure S1*.**

### The expression of ACE2 and TMPRSS2 in normal tissues

In the FANTOM5, HPA, and GTEx databases, ACE2 mRNA and its protein expression were concentrated primarily in the small intestine, testis, kidney, and gallbladder (***Figure 1A***). The mRNA expression of TMPRSS2 showed a nearly fourfold increase in the prostate, stomach, and small intestine. However, the TMPRSS2 protein expression was highest in the kidney (***Figure 1B***). The expression of ACE2 and TMPRSS2 mRNAs were high in the testis, kidney, and colon (***Figure 1CD***) in GSE3526 and GSE8124. There also was high mRNA expression of ACE2 in the kidney, placenta, bone marrow, and testis, which also was confirmed by TCGA. Briefly, the results demonstrated relatively higher expression levels of ACE2 and TMPRSS2 in testis and kidney when the evidence from multiple databases was considered.

**Figure 1.**
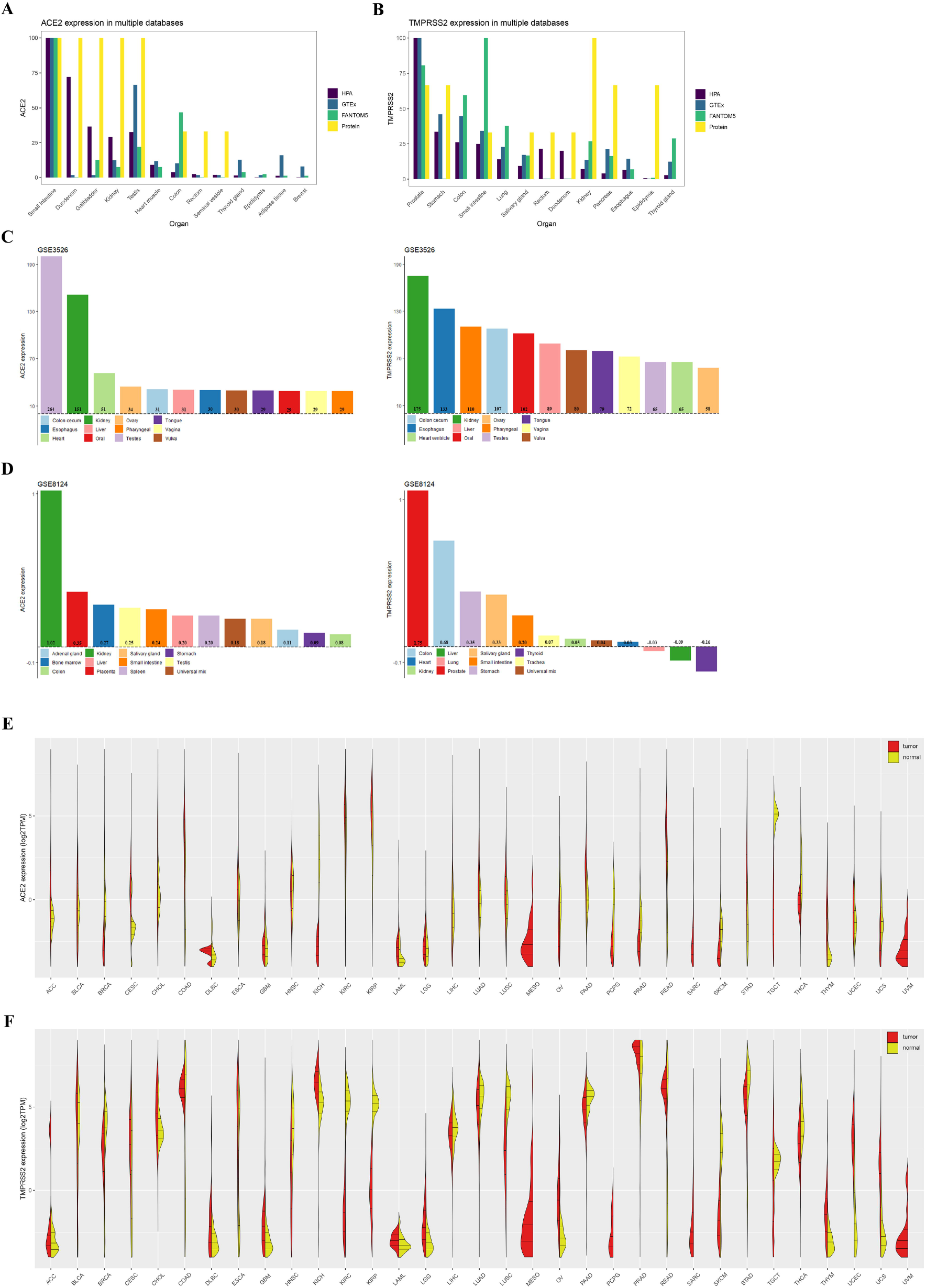
The expression of ACE2 and TMPRSS2 in tissue. **Notes: A**, The mRNA and protein expression pattern of ACE2 in HPA, GTEx and FANTOM5; **B**, The mRNA and protein expression pattern of TMPRSS2 in HPA, GTEx and FANTOM5; **C**, The mRNA expression pattern of ACE2 and TMPRSS2 in GSE3526; **D**, The mRNA expression pattern of ACE2 and TMPRSS2 in GSE8124; **E**, The mRNA expression pattern in TCGA; **F**, The mRNA expression pattern of TMPRSS2 in TCGA.

### ACE2 and TMPRSS2 expression patterns in specific patient populations

Compared with the control group, patients with pulmonary arterial hypertension, chronic kidney disease, and diabetic nephropathy as well as smokers exhibited higher expression levels of ACE2 in their affected tissues (kidneys or lungs) (***Figure 2A-D***). Moreover, there was a significant increase in ACE2 in digestive system tumor tissues (stomach and colon) and lungs (***Figure 2E-F, Figure 1E-F***).

**Figure 2.**
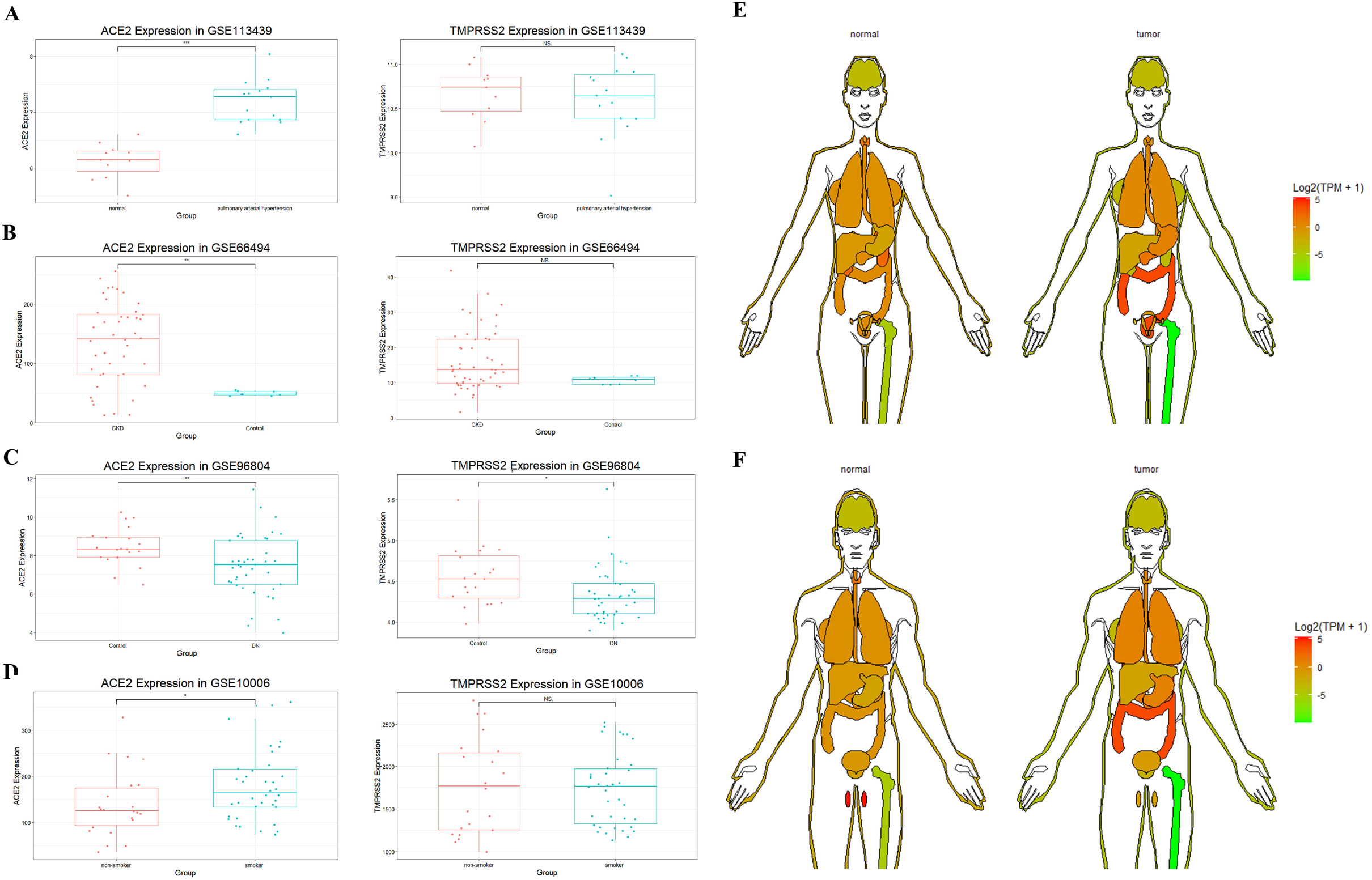
ACE2 and TMPRSS2 expression patterns in specific patient populations. **Notes: A**, ACE2 and TMPRSS2 expression patterns in GSE113439; **B**, ACE2 and TMPRSS2 expression patterns in GSE66494; **C**, ACE2 and TMPRSS2 expression patterns in GSE96804; **D**, ACE2 and TMPRSS2 expression patterns in GSE10006. **E**, ACE2 and TMPRSS2 expression patterns in female. **F**, ACE2 and TMPRSS2 expression patterns in male.

### Multiple sequence alignment - ACE2

To explore the possible causes of animal susceptibility to SARS-CoV-2, we performed a multiple species-sequence alignment for ACE2, which is a key molecule in the process of COVID-19 infection. The results are presented in ***Table 2*** and ***Figure 3***. It appeared that raccoon dogs (distance = 0.142), tigers (distance = 0.144), dogs (distance = 0.144), cats (distance = 0.146), raccoons (distance = 0.154), domestic ferrets (distance = 0.160), masked palm civets (distance = 0.164), domestic cattle (distance = 0.166), and wild boars (distance = 0.172) have an ACE2 sequence that is similar to humans.

**Figure 3.**
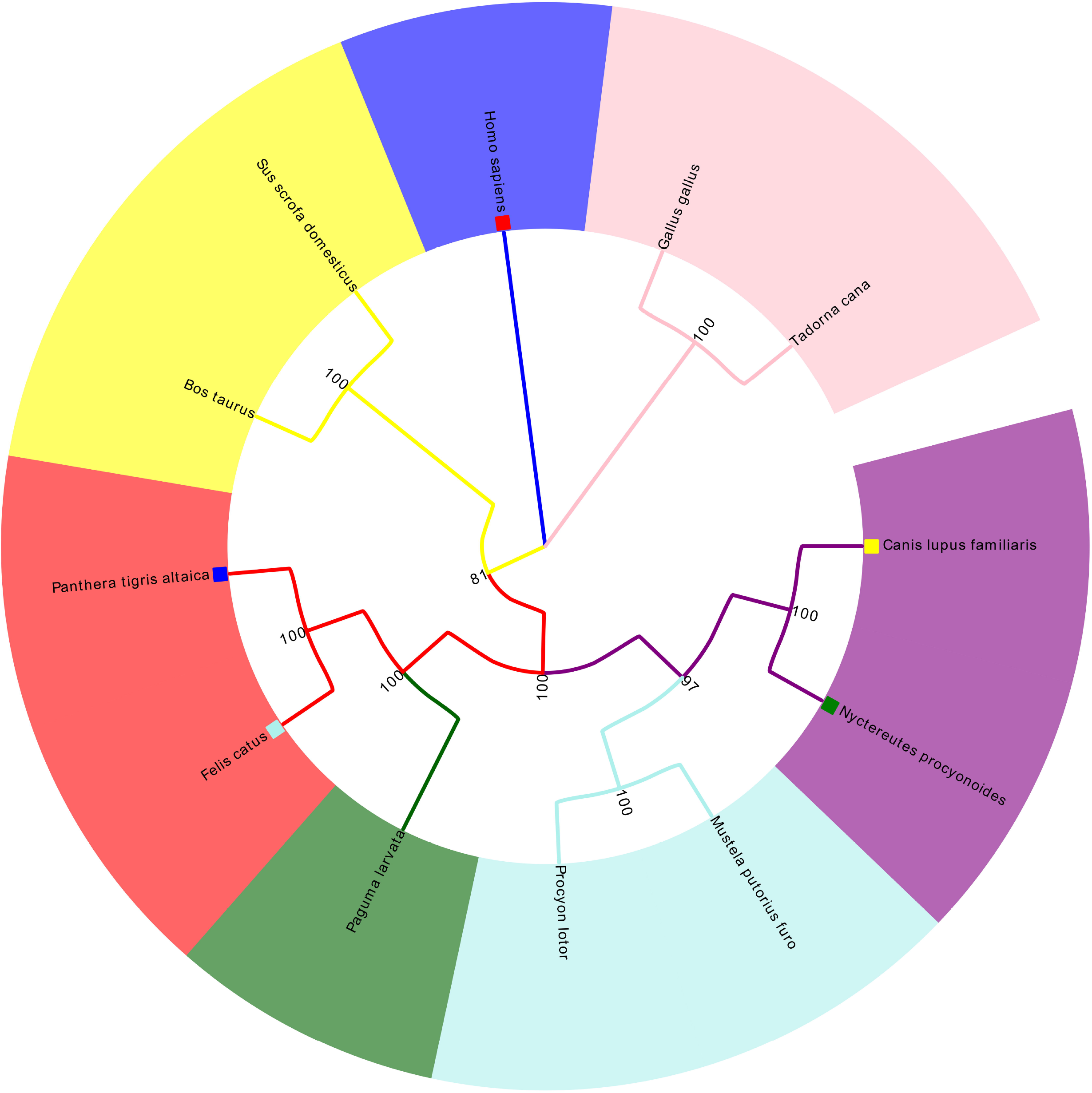
Multiple sequence alignment of ACE2

**Table 2.** The pairwise distance of ACE2 in different species.

|  | 1 | 2 | 3 | 4 | 5 | 6 | 7 | 8 | 9 | 10 | 11 | 12 |
| --- | --- | --- | --- | --- | --- | --- | --- | --- | --- | --- | --- | --- |
| *Homo sapiens* | - |  |  |  |  |  |  |  |  |  |  |  |
| *Tadorna cana* | 0.434 |  |  |  |  |  |  |  |  |  |  |  |
| *Felis catus* | 0.146 | 0.421 |  |  |  |  |  |  |  |  |  |  |
| *Mustela putorius furo* | 0.160 | 0.418 | 0.093 |  |  |  |  |  |  |  |  |  |
| *Canis lupus familiar* | 0.144 | 0.426 | 0.078 | 0.082 |  |  |  |  |  |  |  |  |
| *Gallus gallus* | 0.427 | 0.122 | 0.422 | 0.416 | 0.427 |  |  |  |  |  |  |  |
| *Sus scrofa* | 0.172 | 0.438 | 0.140 | 0.152 | 0.150 | 0.440 |  |  |  |  |  |  |
| *Procyon lotor* | 0.154 | 0.428 | 0.093 | 0.063 | 0.078 | 0.422 | 0.161 |  |  |  |  |  |
| *Paguma larvata* | 0.164 | 0.419 | 0.060 | 0.108 | 0.097 | 0.425 | 0.159 | 0.108 |  |  |  |  |
| *Panthera tigris altaica* | 0.144 | 0.424 | 0.010 | 0.091 | 0.077 | 0.421 | 0.143 | 0.091 | 0.060 |  |  |  |
| *Bos taurus* | 0.166 | 0.440 | 0.141 | 0.152 | 0.139 | 0.437 | 0.121 | 0.148 | 0.152 | 0.138 |  |  |
| *Nyctereutes procyonoides* | 0.142 | 0.423 | 0.075 | 0.079 | 0.009 | 0.424 | 0.150 | 0.077 | 0.094 | 0.075 | 0.138 | - |

**Abbreviations:** Homo sapiens, human; Gallus gallus, domestic chicks; Tadorna cana, waterfowl; cats (Felis catus), Mustela Putorius furo, domestic ferrets; Sus scrofa, wild boars; Canis lupus familiar, dogs; Procyon lotor, raccoon; Paguma larvata, masked palm civets; Panthera tigris altaica, tigers; Bos taurus taurus, domestic cattle; Nyctereutes procyonoides, raccoon dogs.

### Cell-specific expression of ACE2 and cytokine receptors in testis and kidney

To determine the cell-specific expression of ACE2, we analyzed transcriptome data from published single-cell RNA-sequencing studies to decipher the expression pattern of ACE2 in the testis and kidney. ACE2 was highly enriched in the epithelium of renal proximal tubules in the human kidney (***Figure 4A-E***). Considering the critical role of cytokines in disease pathogenesis, we established a signature of cytokine receptors that included CSF1R, CSF2RA, CSF2RB, FGFR2, IFNGR1, IFNGR2, IL6R, IL6ST, IL7R, IL10RA, IL10RB, IL17RA, and NGFR. As displayed in ***Figure 4C-E***, IL6ST, IFNGR1, and IFNGR2 were elevated in various cells in the human kidney, and especially in endothelial and fenestrated endothelial cells. In the testis, from GSE124263, ACE2 exhibited significantly higher expression in gametocytes (***Figure 5A***). Furthermore, from the single-cell RNA sequencing data included in the Human Cell Landscape (Testis; Guo), IL6ST and IFNGR1 expression were notably higher in spermatogonial stem cells, differentiating spermatogonia, Leydig cells, late primary spermatocytes, Sertoli cells, myoid cells, endothelial cells, and macrophages. More than half of the cytokine receptors (IL6ST, IFNGR1, IFNGR2, CSF1R, CSF2RA, IL17RA, IL10RB, and IL10RA) were highly expressed in macrophages (***Figure 5B***).

**Figure 4.**
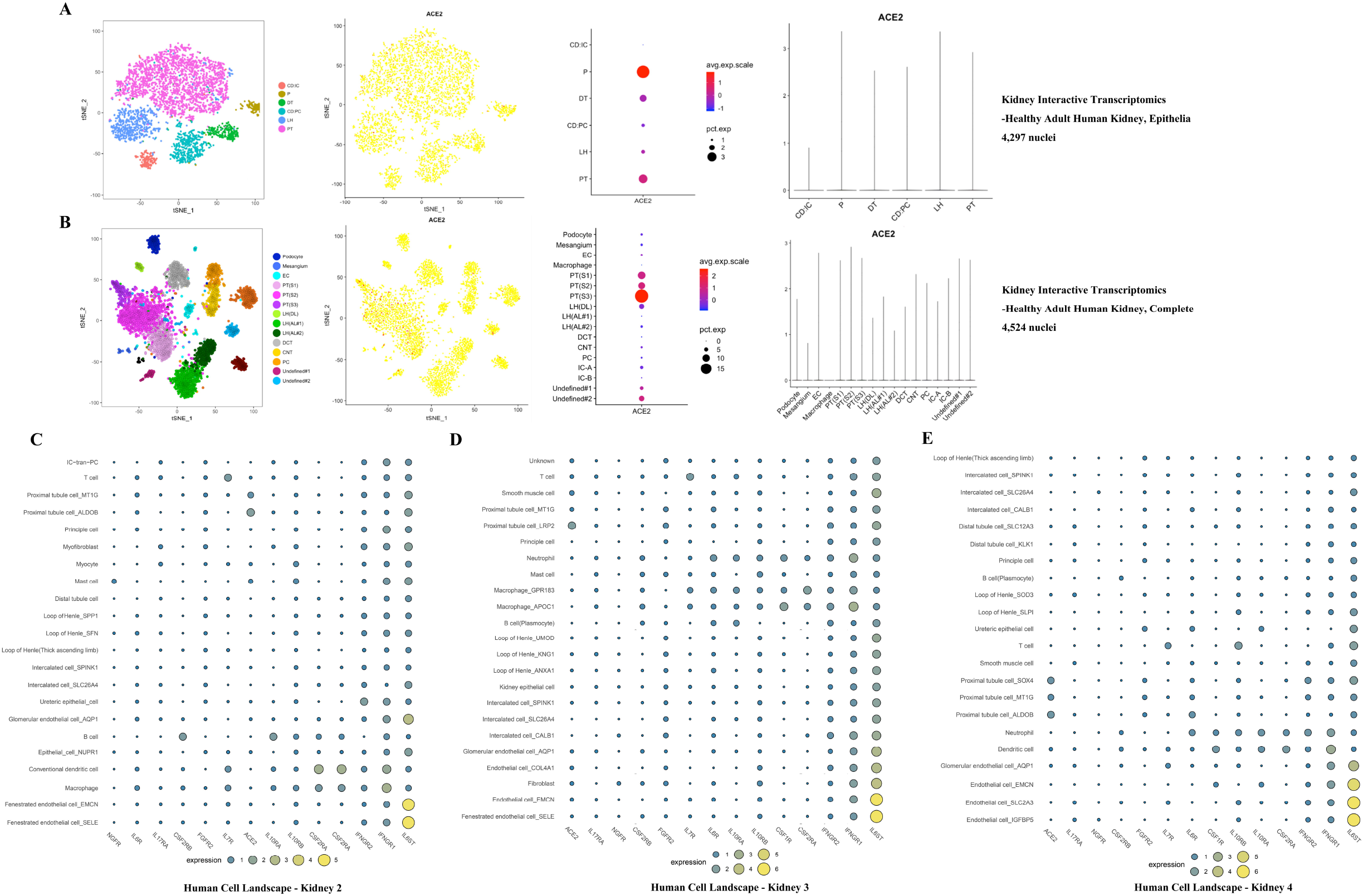
Cell-specific expression of ACE2 and cytokine receptors in kidney. **Notes: A**, Cell-specific expression of ACE2 from http://humphreyslab.com/SingleCell/ (Healthy adult human kidney, epithelia); **B**, Cell-specific expression of ACE2 from http://humphreyslab.com/SingleCell/ (Healthy adult human kidney, complete); **C**, Cell-specific expression of ACE2 and cytokine receptors from Human Cell Landscape Kidney2 (http://bis.zju.edu.cn/HCL/search.html); **D**, Cell-specific expression of ACE2 and cytokine receptors from Human Cell Landscape Kidney3 (http://bis.zju.edu.cn/HCL/search.html; **E**, Cell-specific expression of ACE2 and cytokine receptors from Human Cell Landscape Kidney4 (http://bis.zju.edu.cn/HCL/search.html

**Figure 5.**
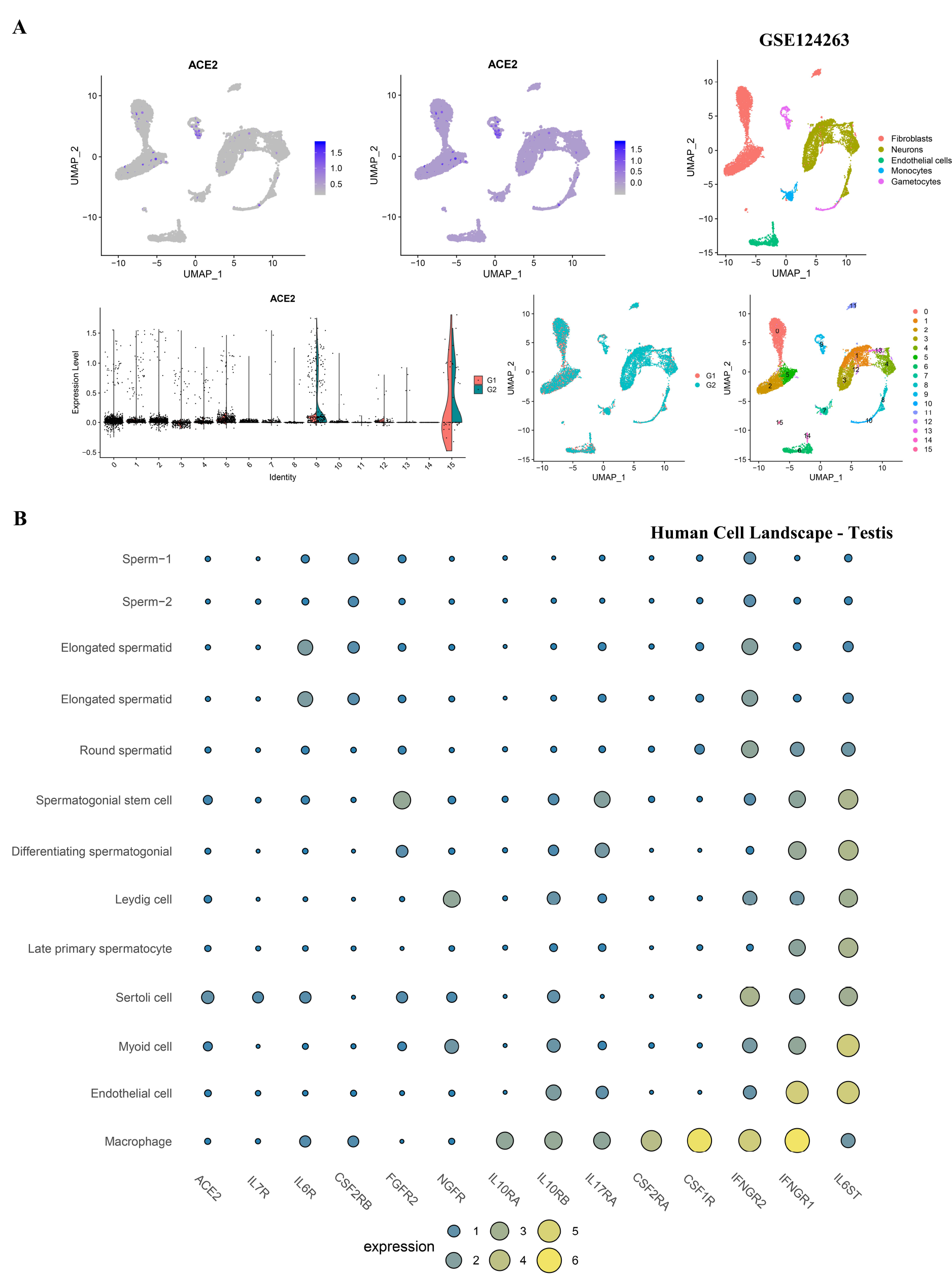
Cell-specific expression of ACE2 and cytokine receptors in testis. **Notes: A**, Cell-specific expression of ACE2 in GSE124263; **B**, Cell-specific expression of ACE2 and cytokine receptors from Human Cell Landscape http://bis.zju.edu.cn/HCL/search.html.

### The biological relevance of the ACE2-specific signature

To speculate on the function of ACE2 in kidney proximal tubules, we generated an ACE2-specific signature from ACE2-positive versus negative expression in kidney proximal tubule cells (***Figure 6A***). The up-regulated functions of ACE2-positive cells were associated with autophagy regulation (GO-bp: regulation of autophagy, negative regulation of autophagy, a process utilizing autophagy mechanism, and others), immunity (GO-bp: myeloid leukocyte mediated immunity, activation of the innate immune response, and others), virus (GO-bp: viral life cycle, positive regulation of viral genome replication, and others), apoptosis (GO-bp: cellular component disassembly involved in the execution phase of apoptosis), and renal physiology (negative regulation of blood vessel diameter and renal water homeostasis). The down-regulated pathways were associated with immunity (GO-bp: macrophage migration, granulocyte migration, and neutrophil migration).

**Figure 6.**
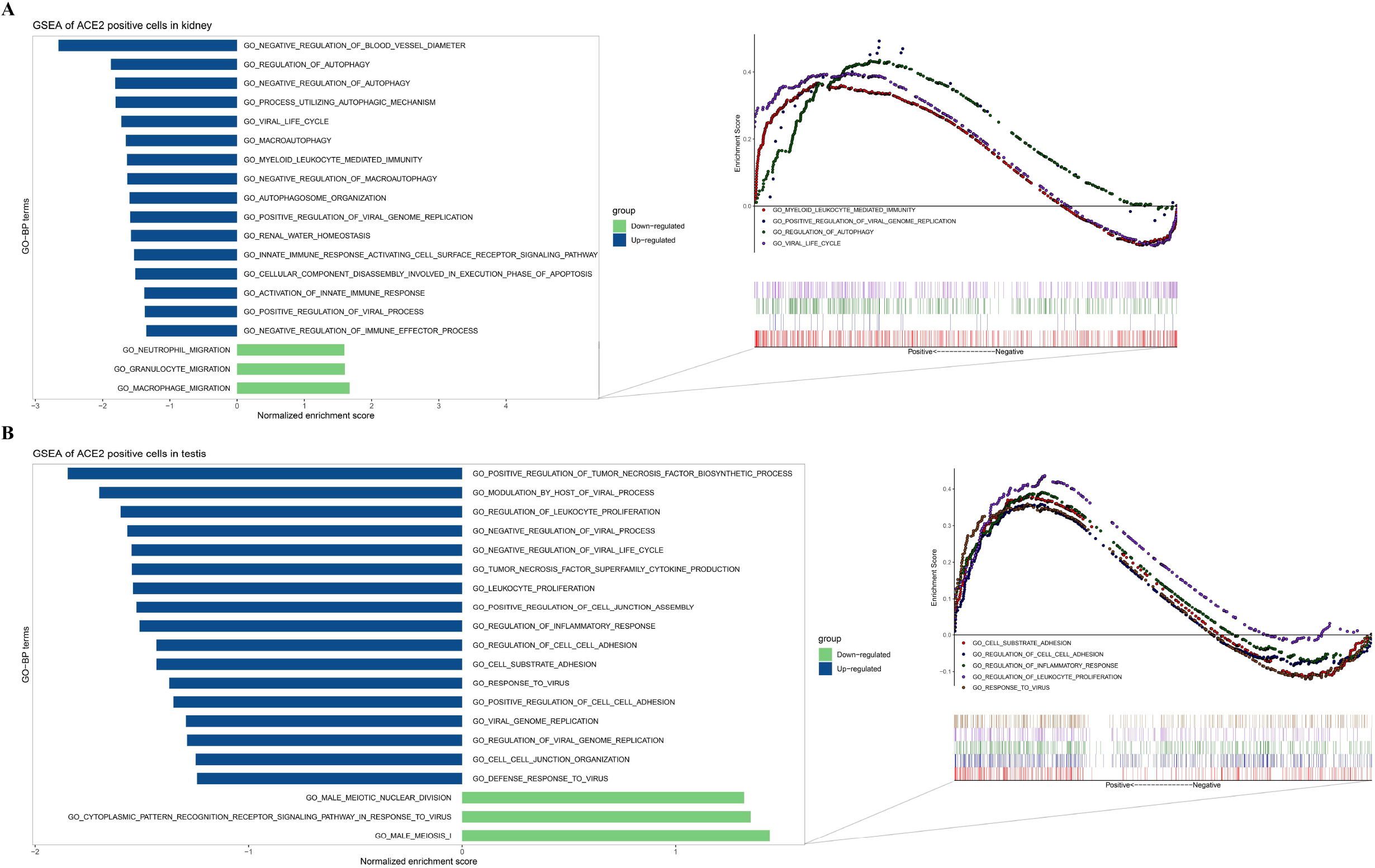
GSEA of ACE2-positive cells in kidney and testis. **Notes: A**, GSEA of ACE2-positive cells in kidney (kidney proximal tubule cells); **B**, GSEA of ACE2-positive cells in testis (spermatogonial stem cells, Leydig cells, Sertoli cells, and myoid cells).

In the testis, we compared the characteristics of ACE2 expression in spermatogonial stem cells, Leydig cells, Sertoli cells, and myoid cells (***Figure 6B***). The up-regulated GO terms in ACE2 positive cells were enriched in cell-cell interactions (GO-bp: regulation of cell-cell adhesion, cell substrate adhesion, positive regulation of cell-cell adhesion, cell-cell junction organization and others), virus (GO-bp: regulation of viral genome replication, response to the virus, viral genome replication and others), and immunity (GO-bp: regulation of leukocyte proliferation, leukocyte proliferation, and others). The down-regulated expression correlated with male reproduction (GO-bp: male meiosis I and male meiotic nuclear division) and response to the virus (GO-bp: cytoplasmic pattern recognition receptor signaling pathway in response to the virus).

## Discussion

With the recent COVID-19 pandemic, it has become apparent that the genitourinary system may be at high risk for injury, which could consequently lead to a range of adverse complications. Thus, this study focused on whether SARS-CoV-2 could directly impair genitourinary organs and what manifestations were observed in the target organs, for example, kidney and testis.

Based on the RNA sequence data from normal tissues in multiple databases (HPA, GTEx, FANTOM5, TCGA, and GEO), ACE2 and TMPRSS2 were highly expressed in kidney and testis. Meanwhile, positive SARS-CoV-2 RNA has been found in the urine sediments from some severely ill patients with urinary tract-related complications^34^. It is tempting to speculate that such complications might be mediated by the entry of SARS-CoV-2 into the kidney via ACE2 and TMPRSS2. Up to now, little direct evidence has been demonstrated for injury to the testis by SARS-CoV-2. However, one study analyzed autopsy specimens of testis from six patients who died from SARS and reported that the SARS virus could induce orchitis, but no positive ISH staining was detected in the testis^35^. Another study reported that SARS-CoV-2 transmission through seminal fluid was hard to achieve and that almost no virus RNA was found in testis cells by qRT-PCR^16^. However, it must be noted that a low detection rate of COVID-19 using PCR was a limitation in these studies. Notably, six of 34 infected patients (19%) experienced scrotal discomfort, which suggested possible testicular damage, although the mechanism was not clear^16^.

Given the possible role of cytokines (especially IL6) in COVID-19 illness, we established a signature that included ACE2 and various cytokine receptors. By performing cell-specific analysis with our signature in the kidney, we found that ACE2 was remarkable higher in proximal tubule cells, and IL6ST (the main receptor for IL6) was significantly enriched in fenestrated and glomerular endothelial cells. From the results of the histopathological examinations conducted by Su et al. on 26 autopsy kidney samples from COVID-19 patients, diffuse injury was observed in proximal tubules and included the loss of the brush border, non-isometric vacuolar degeneration, and even marked necrosis^36^. Besides, nine samples showed increased serum creatinine and (or) the appearance of proteinuria. Our results might partially explain this phenomenon. First, with higher ACE2 expression in proximal tubule cells, the mutual interaction of ACE2 and SARS-CoV-2 may promote virus entry into the cells. Following cell entry, the virus would cause tissue damage via direct injury due to the viral cytotoxicity^37^. Kidney proximal tubule cells with ACE2-positive expression showed some relevant biological events for the mechanisms of injury, including virus regulation, autophagy regulation, and immune and kidney water homeostasis pathways by GSEA. Secondly, the elevated cytokines in infected patients could result in aberrant immune responses and local damage via binding with cytokine receptors in fenestrated and glomerular endothelial cells^38^. This damage would impair the function of the renal glomerular filtration barrier and could feasibly account for the observed abnormalities in serum creatinine and proteinuria.

In the testis, ACE2 is predominantly expressed in gametocytes, Sertoli cells, and spermatogonial stem cells. While this finding suggested that the testis was a high-risk organ that was vulnerable to SARS-CoV-2 infection, direct entry of the virus into testis would be difficult due to the presence of the blood-testis barrier. One study stated that SARS-CoV-2 coronavirus was detected in the semen of male^19^. Despite this, orchitis was reported as a complication of SARS^35^. More importantly, scrotal discomfort in COVID-19 patients has been reported, indicating potential testis injury^16^. From our results, the high expression of multiple cytokine receptors was discovered in testis macrophages, which may disturb their normal functional status, especially in the high cytokine environment of severe COVID-19 patients. The imbalance of macrophages and cytokines is instrumental in the inflammatory response, which could disrupt the functions of Leydig cells, damage the blood-testis barrier, and directly destroy the seminiferous epithelium^35,39^. Meanwhile, the production of inflammatory factors may activate the autoimmune response and auto-antibody development within the tubules, further triggering a secondary autoimmune response^40^. Evidence from a rodent model of autoimmune orchitis demonstrated that increased IL6 and its receptor could result in testicular inflammation and germ cell apoptosis, and ultimately, was involved in the pathogenesis of autoimmune orchitis^41^. Considering the high level of IL6 in severe COVID-19 patients and that IL6 receptor was enriched in various testicular cells, this might be a reason for the potential orchitis caused by SARS-CoV-2 infection. As indicated by GSEA, ACE2-positive cells in the testis might tend to express the genes implicated in the biological process of the virus. Though the specific mechanism is not clear, this finding might provide insight into the testicular damage caused by SARS-CoV-2.

Some studies have reported animal susceptibility to SARS-CoV-2^42,43^. Therefore, we conducted an ACE2 sequence alignment in multiple species, which suggested that felidae and canids (cats, tigers, dogs, and raccoon dogs) shared high sequence similarity of ACE2 with humans. This might be a possible explanation for the fact that seven tigers in the Bronx zoo (New York, USA) were infected with COVID-19 by an asymptomatic zookeeper infected with COVID-19 (https://www.cnn.com/2020/04/05/us/tiger-coronavirus-new-york-trnd/index.html). Also, ACE2 is highly expressed in the kidneys among people with chronic kidney diseases, and diabetic nephropathy, as well as by airway epithelial cells among smokers and lung tissue among people with pulmonary arterial hypertension. These results suggest that the ACE2 expression increases in the corresponding organs of specific populations with underlying diseases, resulting in a higher risk of infection. Thus, particular attention should be given to related symptoms in specific populations.

In conclusion, based on multi-access bioinformatic analyses, this study provided new insights for the impairment of the kidney and testis by SARS-CoV-2 and the possible mechanisms of pathogenesis. These findings indicated that the function of the genitourinary system should be carefully assessed in male patients that recover from COVID-19, particularly in patients with fertility concerns.

## Acknowledgements

This work was supported by the Jiangsu Province “Six Talent Peaks Project” (WSN-011), by the National Natural Science Foundation of China (grant number 81672531, 81972386).

## Disclosure

The author reports no conflicts of interest in this work.

---

**Figure S1.** The visualization map of cumulative confirmed COVID-19 cases.

## Notes

### Competing Interest Statement

The authors have declared no competing interest.

