## Supplementary figures and images for "Multiple expression assessments of ACE2 and TMPRSS2 SARS-CoV-2 entry molecules in the urinary tract and their associations with clinical manifestations of COVID-19"

### Supplemental Figure S1

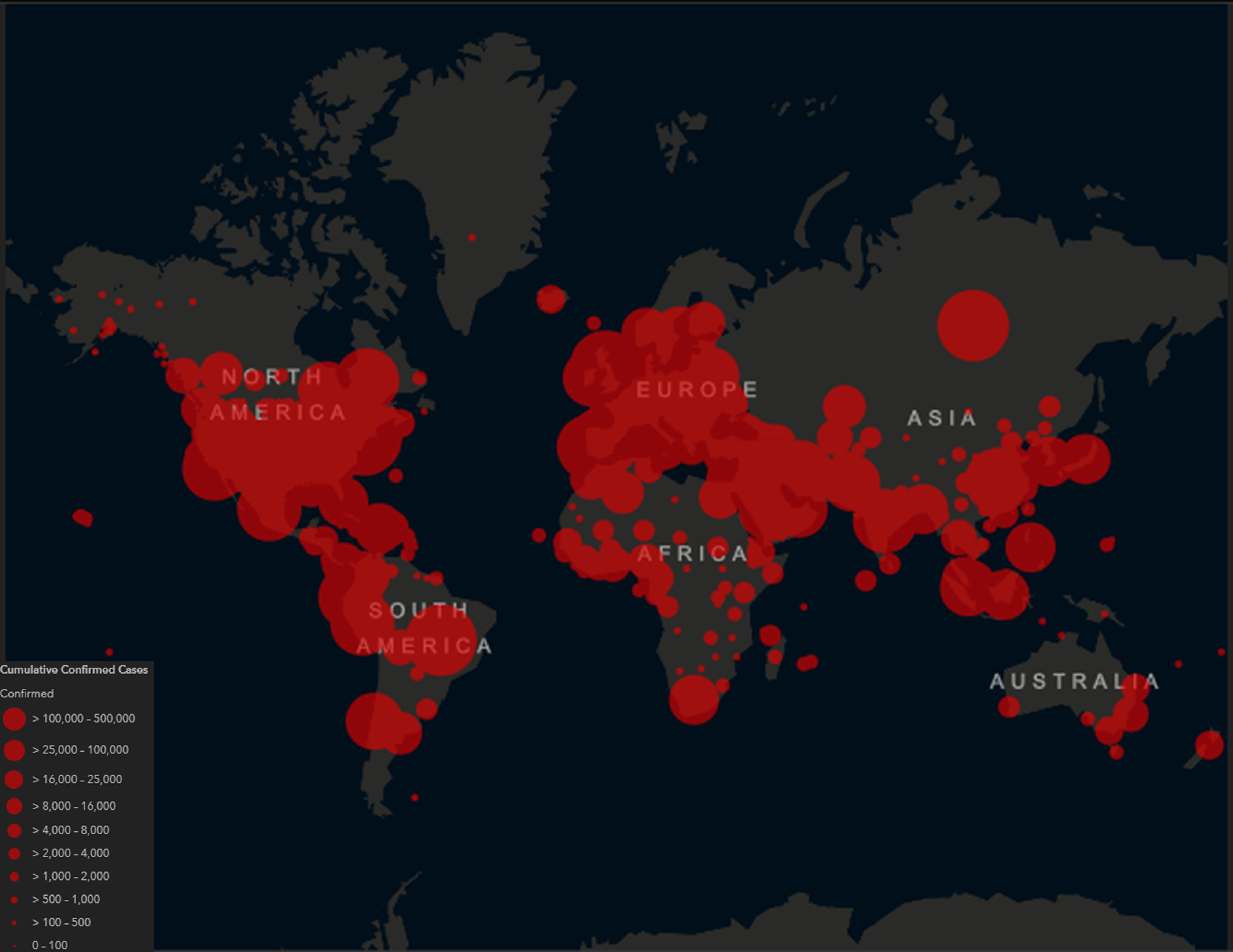
