## Supplemental Table S1 for "Multiple expression assessments of ACE2 and TMPRSS2 SARS-CoV-2 entry molecules in the urinary tract and their associations with clinical manifestations of COVID-19"

| **Table S1. Clinical characteristics of patients infected with SARS-CoV-2** | | | | |
| --- | --- | --- | --- | --- |
|  | Patients at China | | Patients outside of China | |
|  | n=601 | % | n=1310 | % |
| Age (years) |  |  |  |  |
| <20 | 20 | 3.3 | 46 | 3.5 |
| 20-29 | 72 | 11.9 | 64 | 4.9 |
| 30-39 | 121 | 20.1 | 105 | 8.0 |
| 40-49 | 122 | 20.2 | 125 | 9.5 |
| 50-59 | 95 | 15.8 | 170 | 12.9 |
| 60-69 | 101 | 16.8 | 169 | 12.9 |
| 70-79 | 33 | 5.5 | 102 | 7.8 |
| ≥80 | 17 | 2.8 | 68 | 5.2 |
| NM | 16 | 2.7 | 456 | 34.8 |
| Sex |  |  |  |  |
| Male | 311 | 51.7 | 506 | 38.6 |
| Female | 280 | 46.6 | 361 | 27.6 |
| NM | 10 | 1.7 | 443 | 33.8 |
| Symptoms |  |  |  |  |
| Fever | 434 | 72.2 | 568 | 43.3 |
| Cough | 210 | 34.9 | 356 | 27.1 |
| Fatigue | 49 | 8.2 | 23 | 1.8 |
| Muscle pain | 12 | 2.0 | 12 | 0.9 |
| Sore throat | 28 | 4.7 | 86 | 6.6 |
| Headache | 15 | 2.5 | 55 | 4.2 |
| Runny nose | 14 | 2.3 | 32 | 2.4 |
